# Identification and Domestication of a Wild Edible Oyster Mushroom Growing on Loblolly Pine (*Pinus taeda*) in Mississippi

**DOI:** 10.1101/2025.10.23.684254

**Authors:** Frank A. Mrema, Bed Prakash Bhatta, Prachi Bista, Yan Meng, Logan Wiedenfeld, Franklin Chukwuma, Victor Njiti

## Abstract

The oyster mushroom is among the most extensively cultivated edible mushrooms worldwide. These mushrooms typically thrive on decaying broadleaf deciduous trees but are rarely found on conifers. A wild oyster mushroom strain was collected from pine trees in Mississippi. We characterized the strain using both morphological and molecular techniques. Pure cultures of this strain were obtained and utilized for spawn production, cultivation, genomic DNA extraction, and identification. Our research assessed the strain’s capacity to produce extracellular enzymes. The results demonstrated that the strain belonged to the species *Pleurotus ostreatus* (GenBank: PV750134).This fungal strain exhibited a significant ability to colonize lignocellulosic substrates derived from pine, such as logs and sawdust, as evidenced by the substantial production of laccase and cellulase enzymes. Pine logs yielded fruiting bodies approximately 60 days after inoculation. The mushroom strain, ASU_Pre_1-1, shows great potential for producing edible oyster mushrooms using pine wood waste in Mississippi.

## Introduction

The cultivation of edible mushrooms has increased remarkably among small-scale farmers and woodland owners in Mississippi, driven by a recognition of the numerous health benefits and medicinal properties associated with these functional foods (Sreedharan et al., 2025). This trend reflects an increasing appreciation for the nutritional value and unique flavors that mushrooms bring to the table, as well as their potential therapeutic effects. As these farmers tap into the rich potential of their local ecosystems, they are not only enhancing their livelihoods but also contributing to a broader movement toward sustainable and health-conscious food choices. The growing trend in mushroom production is fueled by the increasing demand for fresh, high-quality food, the desire for additional income, and a strong commitment to promoting sustainable food production, thereby improving food security within communities (Mishra & Shankar, 2025).

The most cultivated specialty mushroom crops worldwide are shiitakes (*Lentinula edodes*) and oyster mushrooms (*Pleurotus* spp.). The *Pleurotus* genus is one of the thirteen genera in the Pleurotaceae family, which has over 40 species recorded, and 25 among them are known cultivated oyster species globally (Aditya et al. 2024; Sreedharan et al. 2025; Wang et al. 2024). Of all these edible oyster mushrooms, none is recorded growing on loblolly pine wood substrates because of its resinous extractive contents (Croan, 2003). Conifer wood chips pre-treated with *Ophiostoma pilliferum*, a blue stain fungus in conifers, have been effectively used as substrates for cultivating ten different species of *Pleurotus*, resulting in the successful production of oyster mushrooms with biological efficiencies ranging from 18% to 177% (Croan, 2003). The domestication of wild oyster mushrooms that grow on coniferous trees will enhance our understanding of the biology of mushrooms within the *Pleurotus* genus.

Oyster mushrooms are among the edible fungi that have long been recognized for their therapeutic medicinal properties. These include immunomodulation, antioxidant activity, antiviral and antibacterial effects, antidiabetic benefits, and potential anti-cancer properties. Species of the genus *Pleurotus* contain compounds like β-glucans, which are polysaccharides found in the cell walls of mushrooms. Beta-glucans play several crucial roles, such as activating the human immune system, providing antioxidant and antimicrobial effects, and exhibiting anticancer, antiviral, and antifungal properties (Morris et al., 2017). Additionally, they help regulate cholesterol levels and manage blood glucose levels in the blood. Of all cultivated *Pleurotus* spp., *Pleurotus ostreatus* is the most cultivated oyster mushroom, followed by *Pleurotus pulmonarius* or grey oyster (Jiang et al. 2025; Amirullah et al. 2023).

The cultivation of oyster mushrooms requires certain environmental growth conditions, such as ideal nutrients, temperature, pH, and substrate type, amended with required supplements. Most wild oyster mushrooms typically grow on decaying deciduous broadleaf substrates, which makes them relatively easy to cultivate and identify. In the mushroom cultivation practices adopted by small-scale farmers and woodland owners (SFWOs), wheat straw is often amended with 15 to 20% wheat bran before pasteurization (Oseni et al. 2012). While oyster mushrooms thrive on various lignocellulosic substrates, they are rarely found growing on pine trees in Mississippi.

Consequently, the discovery and characterization of a recently identified oyster mushroom species growing in loblolly pine forests present an opportunity to make use of the significant pine wood waste generated from timber stand improvements. The main purpose of this study was to identify and domesticate a wild edible oyster mushroom strain, ASU_Pre_1-1, collected from loblolly pine (*Pinus taeda*). The morphological characteristics of the mushroom strain, ASU_Pre_1-1, were documented and compared with those in existing literature.

## Materials and Methods

### Pure Culture Isolation

The fresh wild basidiomata were collected from loblolly pine (*Pinus taeda*) forest, in Kemper/Winston County, Mississippi (32º 55’ 39” N 88º 51’ 09” W) kept in a plastic bag in a cooler with ice and processed in the biotech laboratory at Alcorn State University campus for identification and other studies. In the lab, fresh wild basidiomata were used to isolate a pure mycelium culture through the inoculation of fragments of the interior pileus context into petri dishes containing sterile, 20% malt extract agar (MEA) (Merkuri et al., 2016). The voucher specimens were dried and stored for further studies. Morphological features of the collected strain were matched with that of *Pleurotus* species well described in several mycological electronic resources (https://www.mushroomexpert.com/ and https://www.mykoweb.com/).

### Extracellular Enzyme Characterization

A pure culture strain isolated fresh wild basidiomata was investigated for extracellular enzyme production (laccase and cellulase). Except for the media used to test total cellulase in which glucose was not needed, the nutrient solution contained (g/L); NH4NO3 0.6g, K2HPO4, KHPO4 0.5g, MgSO4.7H2O 0.4g and glucose 5.0g. In the Bavendamm reaction test (Etheridge, 1957), the medium solution contained 0.1% guaiacol at pH 5.5 and 25g agar. Assessment was done seven days after inoculation with the isolated strain. For total cellulase, 0.1% cellulose powder was added to the media. All the media were autoclaved at 121°C under 15 psi for 20 minutes. Five days after inoculation of the fungal strain, the plates were flooded with Congo red on the growing mycelium, then cleared within 15 minutes, and examined for cellulase activities (Ogbonna et al. 2018).

### Preparation of the Seed Spawn

Spawn production involved utilizing a blend of grains, including sorghum, millet, and wheat. These grains were thoroughly cleaned three to four times with tap water, followed by a preboiling process for 15 minutes and a soaking period of 12 hours. After draining the water, the grains were enriched with 1% calcium carbonate (CaCO3) and 3% calcium sulfate (CaSO4). The treated grains were then placed in quart jars and autoclaved at 121°C under 15 psi for two hours. Once cooled, the autoclaved jars were inoculated with pure cultures from the wild basidiomata strain and incubated until fully colonized.

### Tested Substrate for the Domestication of Wild Mushrooms

Two lignocellulosic substrates were utilized: *Pinus taeda* sawdust and *P. taeda* logs. Each substrate was thoroughly soaked in water for 12 hours, after which the excess water was drained. The substrates were packed into autoclavable bags, each weighing three pounds, and were amended with 1% calcium carbonate (CaCO3) and 3% calcium sulfate (CaSO4) and autoclaved at 121 ^°^C, 15 psi. After autoclaving, the bags were cooled and inoculated with mother spawn, followed by incubation in the dark for 10 to 15 days. Pinning within the bags began between 24 and 30 days. The colonized substrate was then exposed to enhanced light in an improvised growth chamber, with humidity carefully maintained using a humidifier until the fruiting bodies emerged. *P. taeda* logs (4” diameter x 40” length) were cut from a loblolly pine forest and left for five days. Then holes were drilled in the logs, filled with pine sawdust spawn, and sealed each hole with melted cheese wax. The inoculated logs were placed under the tree shade and irrigated weekly until white mycelium colonized the logs. Spore prints were taken from the successfully harvested mushrooms.

### Molecular Identification of the Strain ASU_Pre_1-1

Around 100 mg of fungal mycelia was collected from a pure culture of the basidiomata growing on MEA petri dishes and ground using liquid nitrogen. The genomic DNA was extracted from the ground mycelium using a commercial DNeasy Plant Mini DNA extraction kit (Qiagen, Hilden, Germany), following the manufacturer’s instructions. The concentration of the DNA was determined using NanoDrop (Thermo Scientific, Delaware, USA). Polymerase chain reaction (PCR) was conducted to amplify the internal transcribed spacer (ITS) region of the fungal genome using ITS-4 and ITS-5 primers (White et al. 1990). The PCR conditions were similar as described in our earlier study (Barnes et al. 2025). The PCR products were purified using the Monarch PCR & DNA Cleanup Kit (New England Biolabs, Ipswich, MA, USA) following the manufacturer’s instructions. The purified PCR products were sent for sequencing to Plasmidsaurus Inc., Louisville, Kentucky, where the sequencing was conducted using the primer-free Oxford Nanopore technology. The obtained ITS nucleotide sequence (655 bp) was trimmed using Geneious Prime® version 2025.0.3, which resulted in the final sequence length of 654 bp. The trimmed sequence was aligned using the National Center for Biotechnology Information (NCBI) nucleotide – Basic Local Alignment Search Tool (BLASTn) website (https://blast.ncbi.nlm.nih.gov/Blast.cgi). The sequence was deposited in the NCBI GenBank with locus ID PV750134.

## Results and Discussion

### Extracellular Enzyme Production of Strain ASU_Pre_1-1

The fresh wild basidiomata collected from the loblolly pine (*Pinus taeda*) forest, in Kemper/Winston County had morphological features that matched perfectly those of *Pleurotus* species well described in several mycological electronic resources (https://www.mushroomexpert.com/ and https://www.mykoweb.com/). The extracellular enzyme production test revealed that the fungal strain ASU_Pre_1-1 was highly positive in laccase production, indicating the ability to produce copious ligninolytic enzymes and can colonize standing host trees (Figure 1). The results from the extracellular enzyme production test indicated that the collected mushroom exhibits a positive reaction for cellulase production. This finding suggests that the mushroom possesses the capability to effectively decompose cellulose and hemicellulose, which are key components of lignocellulosic substrates. The enzyme’s activity is significant as it facilitates the breakdown of complex plant materials, potentially enhancing the mushroom’s ability to exploit various organic matters for growth and nutrient acquisition (Figure 2).

**Figure 1.**
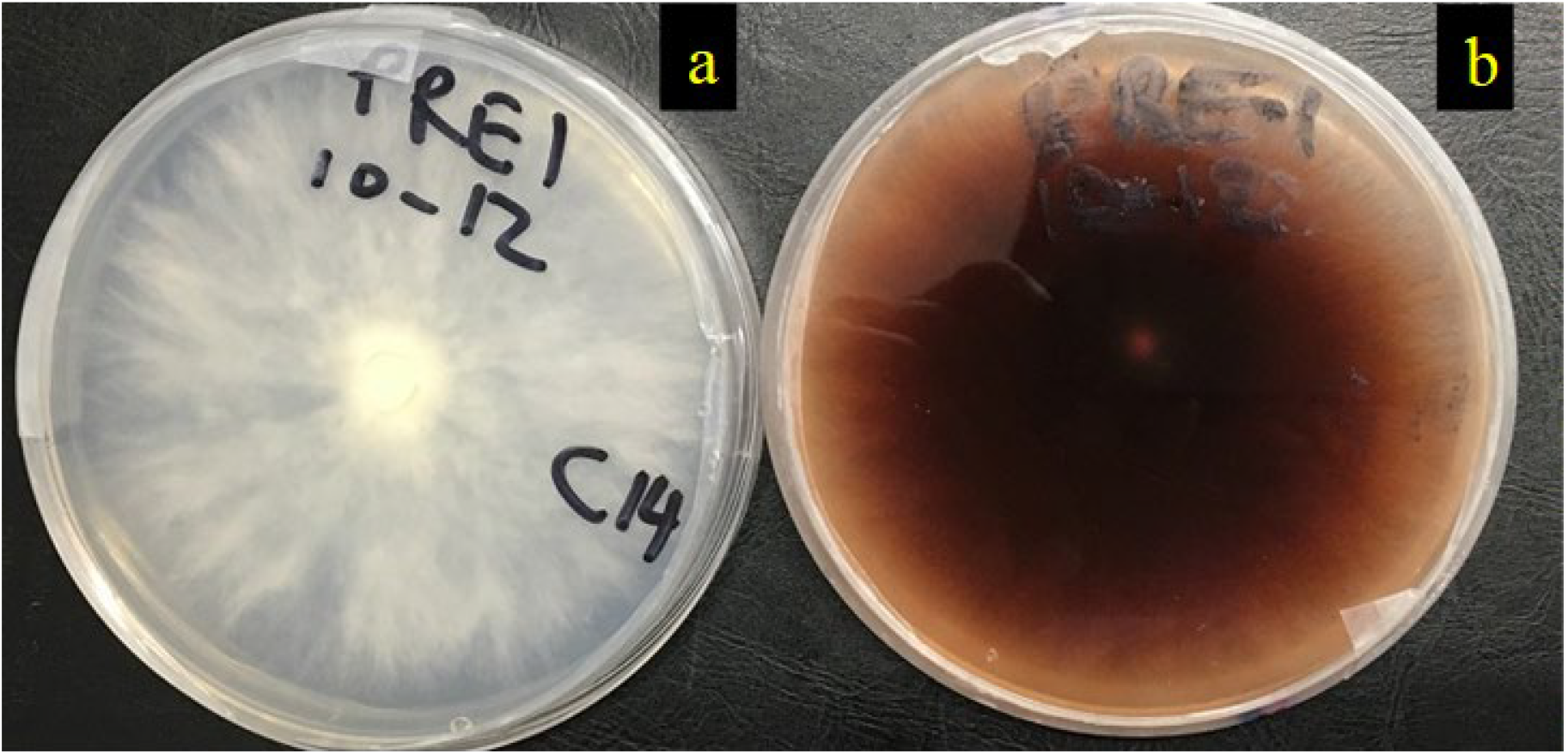
Production of Phenol Oxidase Enzyme in Wild Oyster Mushrooms: (a) Control (Negative) and (b) Treated (Highly Positive) in the Bavendamm reaction test

**Figure 2.**
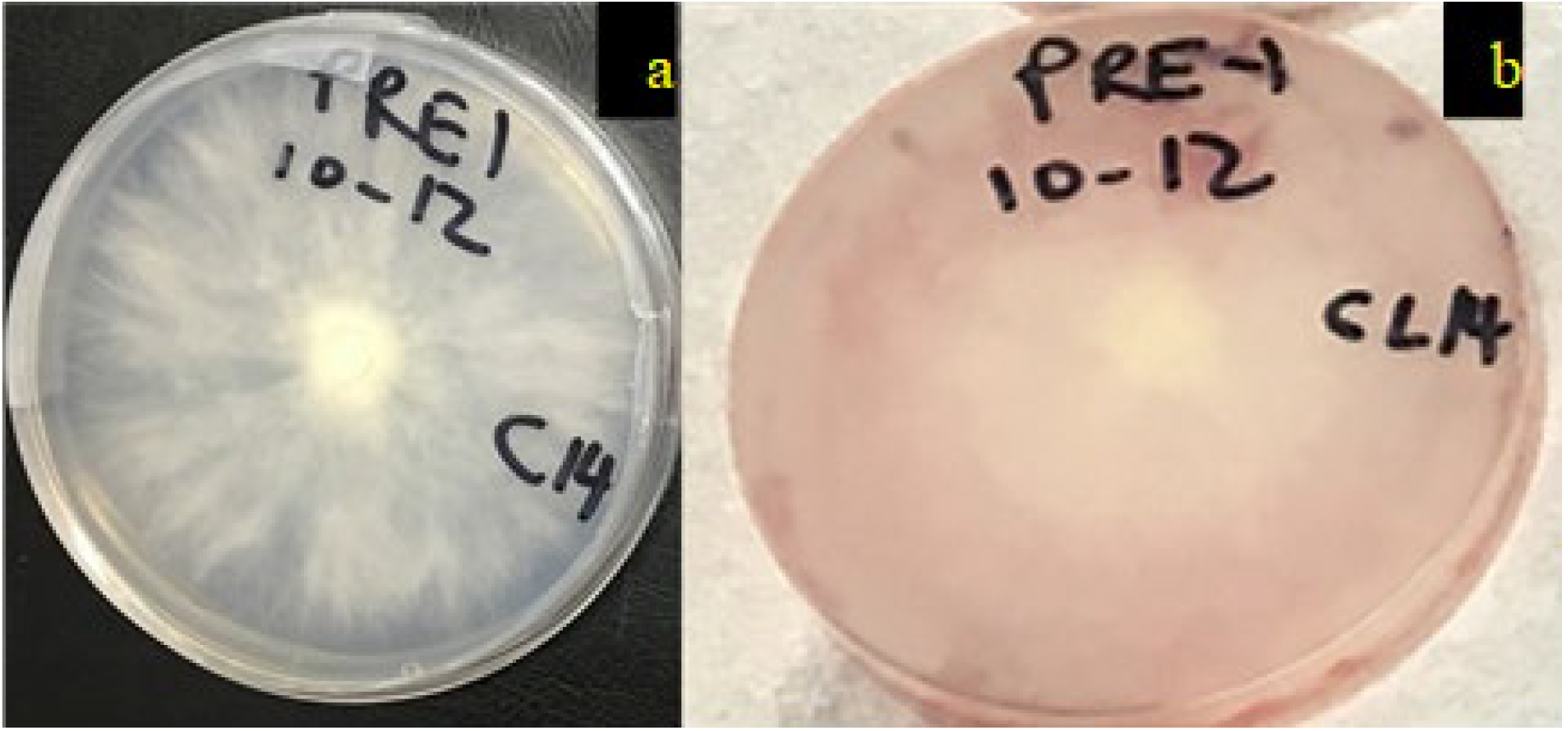
Production of the Cellulase Enzyme in Wild Oyster Mushrooms: (a) Control (Negative in media without cellulose powder), (b) Treatment (Highly Positive in media with cellulose powder)

### Results of Grain Spawn and Pine Sawdust Spawn Preparation

The spawn production incubation period was 16 to 20 days, indicating a successful production of mother spawn. The colonization of the different lignocellulosic substrates was successful. Interestingly, the fungus colonized loblolly pine sawdust 26 (Figure 3).

**Figure 3.**
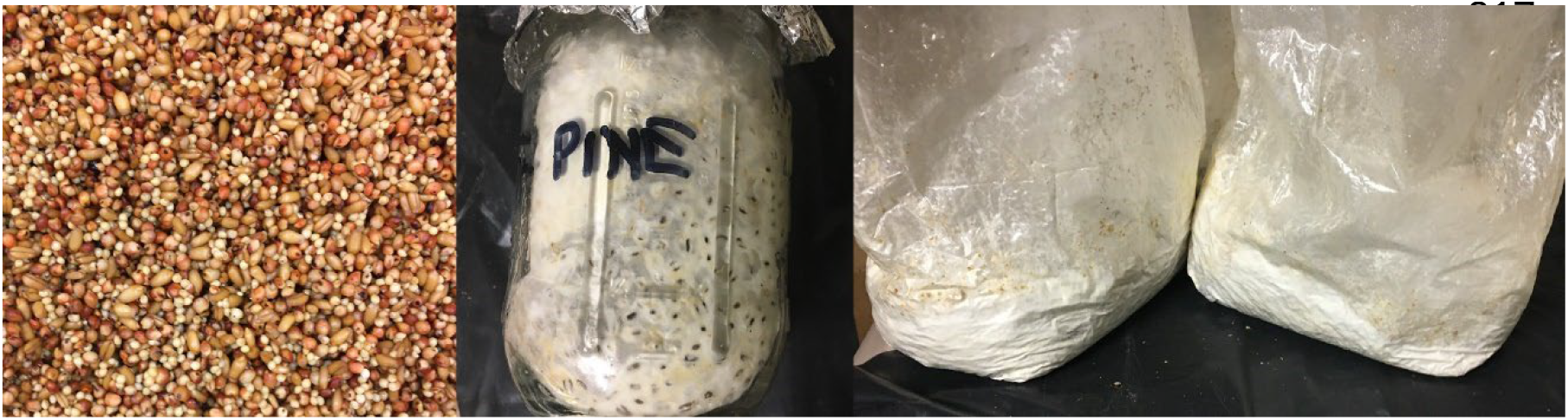
Mycelium Growth of Fungal Strain ASU_Pre_1-1 in Uninoculated Mixed Grains (*left*), Grain Spawn *(middle)*, and Conifer Sawdust Spawn Substrate (*right*).

### Inducing Fruiting Body Formation

Mushroom production was induced on pine sawdust and pine logs. Sawdust spawn was used to inoculate the logs. Figure 4 illustrates the mushrooms that were produced on loblolly pine sawdust. The fruiting bodies produced spores with a lilac color. Figure 5 shows the oyster mushrooms thriving well on pine logs only after 60 days post-sawdust spawn inoculation. Most edible mushrooms colonize the sapwood between the bark. Interestingly, this strain could produce mushrooms even at the end of the logs, suggesting further studies that can explain this growth pattern.

**Figure 4.**
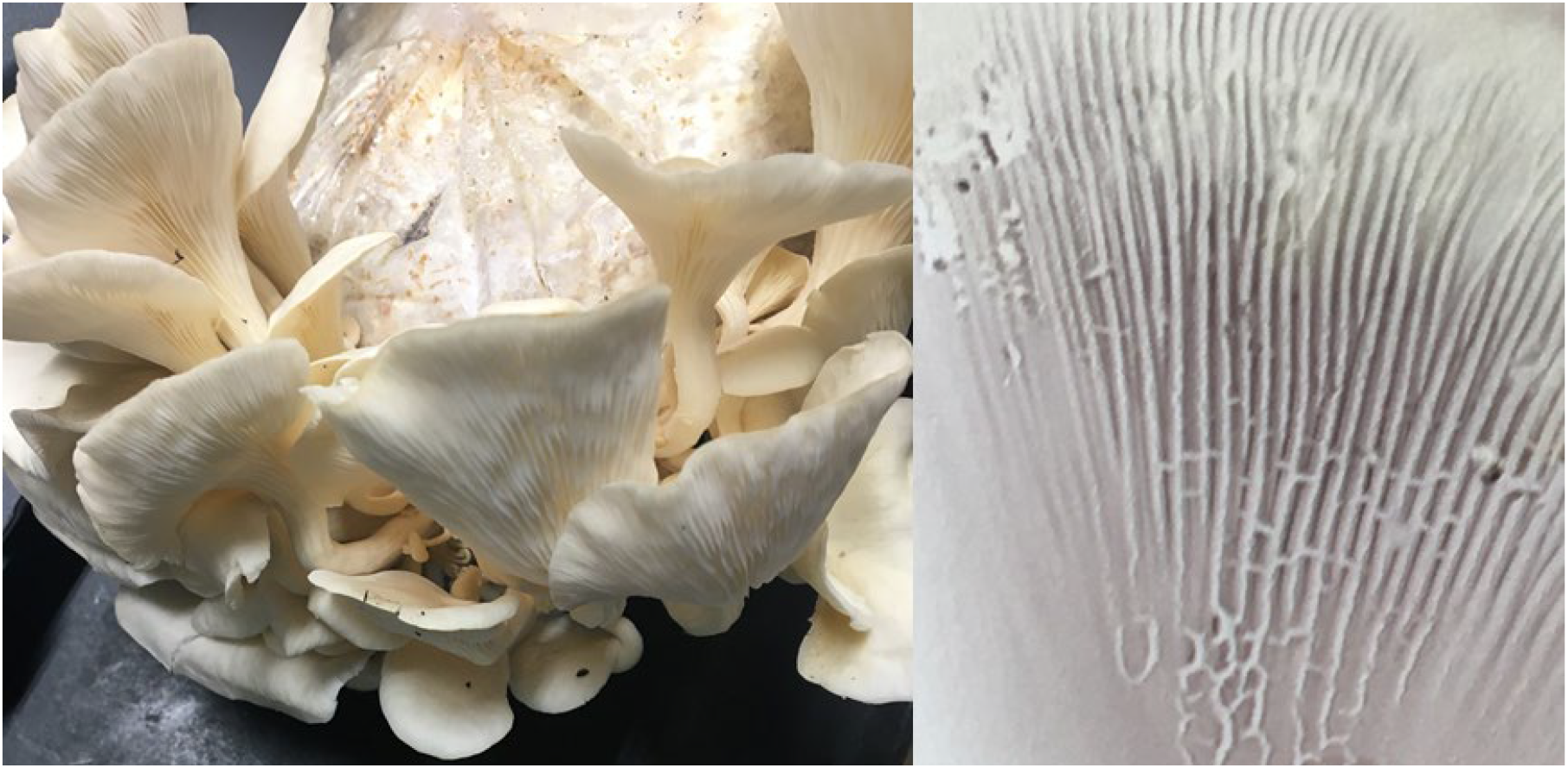
Oyster mushroom (*left*) and spore print (*right*)

**Figure 5.**
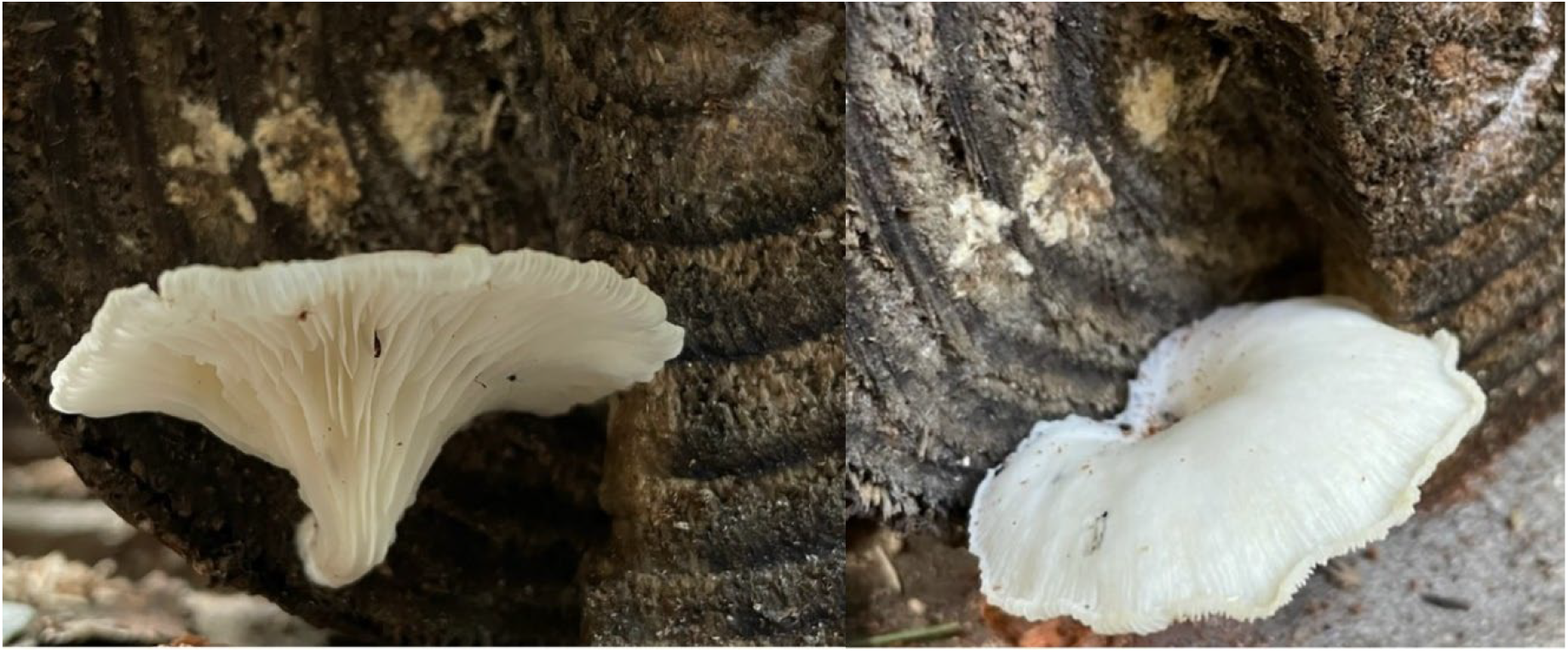
Top and bottom view of *Pleurotus ostreatus* ASU_Pre_1-1 strain thriving at the end of *Pinus taeda* logs

### Molecular Characterization of the Strain ASU_Pre_1-1

The basic local alignment search tool using NCBI BLASTn using both ‘ITS’ and ‘core_nt’ databases confirmed that the strain ASU_Pre_1-1 (PV750134) belonged to *Pleurotus ostreatus* (Table 1).

**Table 1.** Top NCBI BLASTn match for the wild edible oyster mushroom ASU_Pre_1-1.

| GenBank ID of Strain from this Study | Database | Top Match | % Identity | GenBank ID of Matching Strain | Authors of Matching Strain |
| --- | --- | --- | --- | --- | --- |
| PV750134 | ITS | *Pleurotus ostreatus*<br>TENN 53662 | 99.21 | NR_163515 | Matheny et al. 2019 |
|  | core_nt | *Pleurotus ostreatus*<br>strain ICMP 11679 | 100 | MH395969 | Weir et al. 2018 |

### Significance of the wild edible oyster mushroom ASU_Pre_1-1

Wild, edible oyster mushroom strain (ASU_Pre_1-1) from this study can also be useful in utilizing the wood waste generated from the significant damage caused by pine bark beetles (*Dendroctonus frontalis*) in the region (Gomez & Hulcr, 2019). Utilizing pine waste wood as a substrate for mushroom cultivation will enhance the decomposition of organic matter, thereby leading to enrichment of the forest ecosystem while simultaneously producing functional foods (Aditya et al., 2024). This approach offers a profitable means of clearing forests, facilitating sustainable regeneration, and forest establishment. Additionally, this development could yield financial benefits for SFWOs while they await the maturation of timber trees.

## Conclusion

This study demonstrated that the collected basidiomata can successfully colonize loblolly pine woods and produce mushrooms, marking the first evidence in Mississippi that the xylophilous mushroom *Pleurotus ostreatus* thrives on conifer lignocellulosic substrates. The substantial production of extracellular enzymes, including laccase and cellulase, suggests that these enzymes enable the fungus to occupy highly resinous wood substrates and generate basidiomata in such environments. This highlights the necessity for further research in this area.

## Acknowledgments

The authors highly acknowledge the support of the College of Agriculture and Applied Sciences (CAAS) at Alcorn State University. We would like to thank Dr. James O. Garner, Jr. and staff at the Small Farm Incubator Center of Alcorn State University located at Preston, MS. We acknowledge Mr. Leonard Brown of Water Valley, Mississippi and Mr. Michael Maher of McComb, Mississippi for actively participating in growing the oyster mushroom, strain ASU_Pre_1-1 (GenBank: PV750134), on loblolly pine logs.

## Author Contributions

***Conceptualization***, F.M. and B.P.B.; ***Methodology***, F.M., P.B., and B.P.B.; ***Software***, F.M., P.B., and B.P.B.; ***Validation***, F.M. and B.P.B.; ***Formal analysis***, F.M. and B.P.B.; ***Investigation***, F.M. and B.P.B.; ***Resources***, F.M. and B.P.B.; ***Data curation***, F.M. and B.P.B.; ***Writing—original draft preparation***, F.M., P.B., and B.P.B.; ***Writing—review and editing***, Y.M., L.W., F.C., V.N., F.M. and B.P.B.; ***Visualization***, F.M. and B.P.B.; ***Supervision***, F.M. and B.P.B.; ***Project administration***, F.M.; ***Funding acquisition***, F.C., F.M, and Y.M. All authors have read and agreed to the published version of the manuscript.

